# Environmental drivers of earthworm communities along an altitudinal gradient in the French Alps

**DOI:** 10.1101/2022.10.13.512055

**Authors:** Quentin Gabriac, Pierre Ganault, Isabelle Barois, Eduardo Aranda-Delgado, Elisa Cimetière, Jérôme Cortet, Montan Gautier, Mickaël Hedde, Daniel F. Marchán, José Carlos Pimentel Reyes, Alexia Stokes, Thibaud Decaëns

## Abstract

The study of elevational diversity gradients is a central topic in biodiversity research. In this study, we tested for the effect of climate, resource quality and habitat heterogeneity on earthworm communities along an altitudinal gradient and around the treeline in the French Alps. Earthworm communities and environmental properties (i.e. climate, soil properties and vegetation structure and composition) were sampled in six altitudinal stages from 1400 to 2400 m. Results were analysed through multi-table factorial analyses and structural equation modelling. We found average density, biomass and species richness in the range of what is usually reported in comparable ecosystems. We found no monotonic decrease in species richness along the altitudinal gradient, which we explain by the species pool being dominated by taxa with high environmental tolerance and dispersal capacities. Instead, we highlighted the ecotone associated with the treeline as the primary driving factor of earthworm communities: at 1800-2000m altitude, communities were more abundant and diverse, and had a greater variability in body mass. This result was largely explained by the structure and composition of the vegetation, whereas soil and climate appeared to have only indirect effects. Therefore, the treeline effect on earthworm communities can be explained both by the effect of environmental heterogeneity and of trophic resource quality which increases at the ecotone level.

## 1. Introduction

Studying the variations in the diversity and composition of ecological communities along altitudinal gradients has long fascinated early naturalists and ecologists, and still represents nowadays a strong topic in biodiversity research [1,2]. The study of elevational diversity gradients (EDG) has contributed to a better understanding of the links between the environment and the diversity and structure of ecological communities, and is expected to provide insights into the responses of species and ecological systems to ongoing global changes [3]. Understanding EDG is therefore fundamental for establishing conservation and management strategies for mountain ecosystems that are particularly sensitive to global warming and often subject to significant anthropogenic pressures [4].

Most studies of EDG describe a decline of species richness with increasing elevation, but whether this decline is monotonic or if it assumes different shapes depending on the taxa or region being studied has been an important topic of debate [5]. Currently, it is broadly accepted that the most widespread pattern corresponds to a mid-elevation peak of species richness, and that the monotonic variations sometimes described corresponds to studies having considered an incomplete gradient unable to fully cover environmental heterogeneity [6]. Mid elevational peak in species richness has received considerable empirical evidence for different taxa, such as mammals [7,8], birds [9], arthropods [10–13], and vascular plants [14] among others, but was not documented for other organisms such as soil microbes [15]. Different hypotheses have been proposed to explain EDG, most of them involving the effects of elevational gradients in habitat area, in ecosystem biophysical conditions, in the degree of geographical isolation of montane biota and in turnover among zonal communities [1,4,5]. Alternatively, in some cases the observed patterns could result from the spurious outcome of bias in sampling regimes [1].

Another feature of altitudinal gradients that can influence EDG locally is the presence of a treeline, also called timberline, which marks the upper limit in tree distribution and the transition between low elevation closed forest and upper elevation alpine tundra. The study of the dynamics of this ecotone is a major challenge for the conservation of mountain ecosystems, especially in the context of the current global changes which are likely to modify both its functioning and altitudinal position [16–18]. However, studies of ecological communities on both sides of the treeline have so far focused mainly on plant communities, leaving out other functionally important groups of organisms such as soil invertebrates (but see [19,20]). Among them, earthworms deserve special attention because of their capacity, as ecosystem engineers, to modify soil structure and water properties, and to stimulate the decomposition of organic matter and the recycling of nutrients [21]. Paradoxically, there are very few studies to date on the way earthworm communities may vary in structure and diversity along altitudinal gradients and around the treeline.

In addition to the above factors proposed to explain variations in biological diversity along altitudinal gradients, vegetation composition is also likely to strongly influence earthworm communities. Plant species can influence soil detritivorous fauna through the provision of energy and matter via living and dead plant products such as leaf and root litter and rhizodeposition [22]. Deciduous trees in particular are known to produce a litter of higher palatability compared to conifers [22,23], and alpine herbaceous plants also produce a high quality trophic resource compared to trees and shrubs, either through litter or dead roots [24,25]. At comparable altitudes, earthworm abundance and diversity thus tend to decrease with the proportion of conifers in the standing vegetation [22,23], and to be higher in pasture compared to forest ecosystems [26,27]. Along the altitudinal gradient, observable changes in vegetation composition, including variations in the proportion of conifers, as well as the shift to more herbaceous vegetation above the treeline, are therefore likely to be associated with changes in earthworm biomass and diversity.

In this study, we sought to understand the factors controlling earthworm communities along an altitudinal gradient around the treeline in the French Alps. We tested three alternative hypotheses that could explain the variation of earthworm diversity along this elevation gradient: 1) the climate hypotheses, which assumes that earthworm biomass and diversity should decrease with altitude as a result of increasingly adverse temperature conditions [1,28], and that communities should be dominated by large body-sized species at higher altitude as a result of the Bergmann’s rule [29]; 2) the resource quality hypothesis, which predicts an increase in earthworm biomass and body mass with elevation as a result of the shift from gymnosperm stands to alpine grasslands, and the consecutive increase in the quality of litter inputs to the soil food web [30]; 3) the habitat complexity hypothesis, which assumes that community taxonomic and functional diversity will peak at the ecotone level due to higher diversity in microhabitats [8].

## 2. Material and methods

### 2.1. Study sites

The study was carried out in the Belledonne massif in the French Alps (Chamrousse, Isère, France, N45°7’1” E5°53’35”) (Fig. A.1), along a 8 km long altitudinal gradient ranging from 1400 to 2400 m above sea level. It was part of a project whose objective was to explore the relationships between biological communities (vegetation, soil invertebrates and microbial communities) and soil properties along altitudinal gradients in France and Mexico (ECOPICS project, ANR-16-CE03-0009 and CONACYT-2 73659). A dataset comprising detailed information about climatic data, vegetation and soil biological, chemical and physical properties is available at [31].

The geological substrate was composed of Variscan metamorphic rocks and ophiolitic complexes, and soils were classified as Umbrisols and Cambisols [31,32]. Climatic data were estimated over 2004–2014, using the Aurelhy model [33]. Vegetation consisted of mixed and coniferous forests to alpine meadows, with a transition between forest and meadows (i.e. ecotone) at 1800-2000m and a treeline located between 2000 and 2100 m (Fig. A.2).

Along this altitudinal gradient, sampling was carried out at six altitudinal levels spaced every 200 m of elevation from 1400 to 2400 m. Five plots (20 × 20 m) were selected within each altitudinal band, with a lateral distance of at least 100 m between each plot. Plots had a south-west orientation (azimuth of 220 ± 26°) and a mean angle of 17.5 ± 2.3° from the vertical. Areas with visible bedrock or signs of waterlogging were avoided. Plots were selected if they included three species: *Picea abies* (L.) H. Karst (Pinaceae), a tall evergreen tree; *Juniperus communis* L. (Cupressaceae), a prostrate evergreen shrub and *Vaccinium myrtillus* L. (Ericaceae), a small deciduous shrub (if these species were present at that elevational band). These three plant species were selected because of their different growth forms and occupation of different ecological niches along the elevational gradient. *P. abies* was the dominant tree species below the tree line; *J. communis* was abundant locally and *V. myrtillus* was one of the most frequent species above the treeline, as well as being present in all six elevational bands. *P. abies* and *V. myrtillus* are both keystone species [34,35] and although *J. communis* is not classed as a keystone species, its abundance above the treeline makes it an important species that structures plant communities. Therefore, these three species contribute to shaping the structure of the plant communities in which they are present, and we term them ‘structuring species’ [36]. Structuring species were chosen because of their different growth forms and their ecological niche along the elevation gradient. *P. abies* is the dominant tree below the treeline, *J. communis* is abundant in the shrub layer above the treeline, while *V. myrtillus* occurs in the shrub layer along the entire altitudinal gradient. Therefore, the importance of these species suggests that they could have a significant long-term influence on the biotic (i.e. plant – plant and plant – soil interactions) and abiotic (i.e. physical and chemical properties) soil properties along the elevational gradient.

The proportion of the ground covered by trees, shrubs, herbs, and bryophytes, the proportion of mineral elements and bare soil, as well as the relative proportion of the three structuring plant species within their own strata were visually estimated on each 20 × 20 m plot (Fig. A.2). *V. myrtillus* was present at all elevation levels and was more abundant below the treeline (17% vs 2% on average). The relative cover of *P. abies* among trees decreased steadily from 31% at 1400 m to 3% at 2000 m. *J. communis* was present from 1800 to 2400 m with an average relative coverage of 8.5%.

### 2.2. Sampling and description of earthworm communities

Within each 20 × 20 m plot, a 25 × 25 × 15 cm soil block (or monolith) was taken from under one individual of each of the structuring plant species present. The exact location of this monolith was at the limit of the projection on the ground of the canopy of the individual plant. Earthworms were collected by hand sorting each monolith in a large tray, immediately fixed in 100% ethanol, and stored at −21° C. Total sampling varied from 10 to 15 monoliths per altitudinal level depending on vegetation composition.

Adult and sub-adult individuals were identified at species level on the basis of their external morphology [37]. Identification of juveniles, which often do not present the necessary morphological characters, were achieved by sequencing the DNA barcode of the cytochrome c oxidase I (COI) gene [38] through standard protocols [39] implemented at the Canadian Centre for DNA Barcoding (CCDB), University of Guelph, Ontario, Canada. Sequences obtained for juveniles were further compared to those obtained for adult specimens, or alternatively to identified sequences already available in the Barcode of Life Data systems (BOLD; [40]). This enabled the species-level identification of all the unidentifiable specimens for which DNA barcoding was successful [41]. All sequence records, voucher deposition details, specimen data, GPS coordinates and photographs are publicly available within the DS-EWCHAM dataset in BOLD (dx.doi.org/10.5883/DS-EWCHAM).

Community functional properties were described using trait-based ecological categories and species body mass. We used data from Bottinelli et al. [42] to attribute to each species affinity scores regarding the three ecological categories defined by Bouché [37] (Table A.2). These categories can be broadly described as follow [42]: epigeics are surface dwelling species, small sized, light in weight and present pigmented body; anecic live in vertical burrows and are relatively large and pigmented species with flattened tail; endogeics are soil dwelling species deprived of pigmentation, with variable body size. Each individual was then weighed individually after being carefully dried, and body mass values were aggregated to obtain average biomass.

For each plot we calculated the community weighted mean (CWM) of ecological category scores and of body mass, and the community weighted variance (CWV) of body mass using the following formula [43–45]:

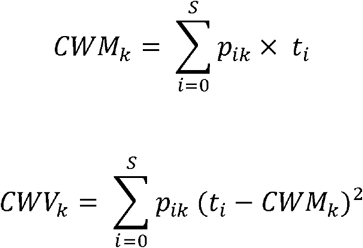

where *S* is the number of species in community *k, p_ik_* is the relative abundance of species *i* in community *k*, and *t_i_* is the mean value taken by trait *t* and for species *i*.

### 2.3. Soil properties

At each monolith location, the depth of litter and organo-mineral layers was recorded on three randomly chosen sides of the pit. Bulk density was measured on a 5 cm diameter and 5.9 cm depth soil core (i.e. 115.89 cm^2^] which was dried at 105° C before being weighed (BD expressed in g of dry soil. cm^−2^). Soil physical and chemical properties were measured at the INRA-Arras (France) laboratory on a sample aliquot taken from each monolith, air dried and sieved at 2 mm. Soil texture was determined on non-decarbonated soil using the Robison method (NF X 31-107) [46]. Water pH was measured using the NF ISO 10390 method (H_2_O/KCl/CaCl_2_). Different soil chemical properties, which are expected to be indirectly related to the quality of litter inputs by the vegetation, were also measured. NaHCO_3_ extractible phosphorus was measured at a pH of 8,5 (NF ISO 11263) using the Olsen et al. method [47]. The cationic exchange capacity (CEC, expressed in cmol. kg-^1^) was estimated in a cobalihexamine solution following the NF X 31-130 method [48]. Total organic carbon and total nitrogen were determined after dry combustion using the NF ISO 10694 and NF ISO 13878 methods, respectively [49]. NH4^+^ and NO_3_^-^ were measured on soil samples conserved at −20 °C after KCl extraction, using colorimetric tests as recommended by ISO 14256-2) (analyses done at US Analyses, Cirad, Montpellier).

### 2.4. Statistical analyses

The overall effect of elevation and vegetation (i.e. structuring plant species under which the monolith was sampled) on earthworm biomass (g of fresh mass, m^−2^), abundance (number of individuals. m^−2^), body mass CWM (g) and body mass CWV were tested using Kruskal-Wallis rank sum test with the ‘kruskal.test’ function of R. Pair wise differences among altitudinal levels and plant species were tested using pairwise Wilcoxon rank sum tests with the ‘pairwise.wilcox.test’ function of R 4.0.3 [50].

Local species richness was calculated as the cumulative number of species observed in the monoliths of a given study plot. To compare the richness of the species pools corresponding to each altitudinal level, the data from the study plots were pooled in order to plot one rarefaction curve (intrapolation and interpolation) for each altitude. This was done using the ‘iNEXT’ package considering the number of individuals as a measure of the sampling effort. We then used the same package to calculate the asymptotic estimator of Chao for each altitudinal level [51–53].

The relative effect of environmental parameters on the structuring of earthworm communities was assessed through a set of factorial analyses. The data were first organised into six tables sharing 30 rows corresponding to the study plots (i.e. five plots in each of the six altitudinal levels). The earthworm data were thus formatted into a table containing species abundance, and a table describing community structure through a set of selected metrics (see Fig. A.4 for a list of species and metrics used in these two tables). The environmental data were organised into a *Clim* table containing climatic variables, a *Soil* table containing soil physicochemical properties, a *Veg1* table containing vegetation structure descriptors and a *Veg2* table containing vegetation compositional variables (see Fig. A.3 for a list of these variables).

The earthworm species abundance table was first analysed using a principal component analysis (PCA) after applying a Hellinger transformation to the abundance data using the ‘deconstand’ function of ‘Vegan’ package [54]. We then used the scores of the study plots on the first two axes of this PCA as two synthetic composition variables named *Com.1* and *Com.2*. These two variables were added to the community structure variables to constitute an *EW structure* table that was further analysed with another PCA. The scores of study plots on the axes of this PCA were extracted and referred to as *Str.1* and *Str.2* in further RDA and path analyses (see Fig. A.4 for a complete description of this analysis).

The main axes of environmental variability (i.e. climate, soil and vegetation) along the environmental gradient were also analysed through four individual PCAs. The scores of the study plots on the significant axes of these PCAs were used as synthetic environmental variables for further analyses. By significant axes we mean those axes of a particular PCA that when combined explained at least 70% of the total variance in the analysed table. We thus retained the first axis of the PCA of climatic variables (*Clim.1*), as well as the first two axes of the other three PCAs (denoted *Soil.1, Soil.2, Veg1.1, Veg1.2, Veg2.1* and *Veg2.2*, respectively). The synthetic interpretation of each of the retained axes is given in Table 1 (see complete interpretation in Appendix 2) All these analyses were done using the ‘ade4’ package for R [55].

**Table 1.**
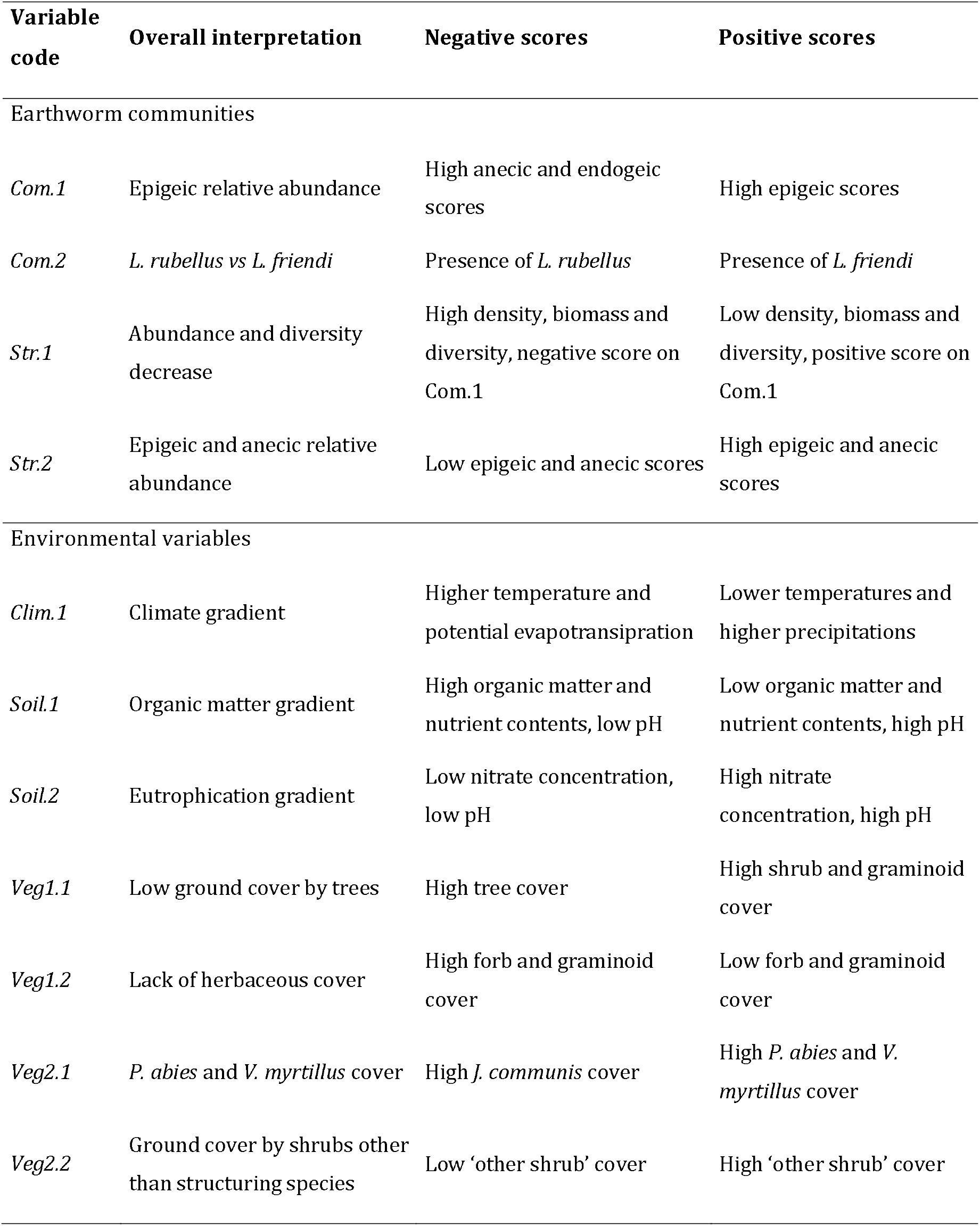
Synthetic interpretation of the axes of the different PCA used in the analyses to describe earthworm communities and environmental variables (see Fig. A.3 for complete description of the analyses).

| Variable code | Overall interpretation | Negative scores | Positive scores |
| --- | --- | --- | --- |
| Earthworm communities |  |  |  |
| *Com.1* | Epigeic relative abundance | High anecic and endogeic scores | High epigeic scores |
| *Com.2* | *L. rubellus* vs *L. friendi* | Presence of *L. rubellus* | Presence of *L. friendi* |
| *Str.1* | Abundance and diversity decrease | High density, biomass and diversity, negative score on *Com.1* | Low density, biomass and diversity, positive score on *Com.1* |
| *Str.2* | Epigeic and anecic relative abundance | Low epigeic and anecic scores | High epigeic and anecic scores |
| Environmental variables |  |  |  |
| *Clim.1* | Climate gradient | Higher temperature and potential evapotranspiration | Lower temperatures and higher precipitations |
| *Soil.1* | Organic matter gradient | High organic matter and nutrient contents, low pH | Low organic matter and nutrient contents, high pH |
| *Soil.2* | Eutrophication gradient | Low nitrate concentration, low pH | High nitrate concentration, high pH |
| *Veg1.1* | Low ground cover by trees | High tree cover | High shrub and graminoid cover |
| *Veg1.2* | Lack of herbaceous cover | High forb and graminoid cover | Low forb and graminoid cover |
| *Veg2.1* | *P. abies* and *V. myrtillus* cover | High *J. communis* cover | High *P. abies* and *V. myrtillus* cover |
| *Veg2.2* | Ground cover by shrubs other than structuring species | Low 'other shrub' cover | High 'other shrub' cover |

We then ran a redundancy analysis (RDA) to figure out the relative importance of climate, soil and vegetation in explaining altitudinal variations in earthworm community structure. This was done using the ‘rda’ function of the ‘Vegan’ package. The community data was the *EW structure* table, and the explicative variables were the plot scores on the retained axes of the PCAs of environmental data. The model formula (‘*EW.structure.table~Clim.1+Soil.1+Soil.2+Veg1.1+Veg1.2+Veg2.1+Veg2.2’*) included all possible interactions among the different explanatory variables. Permutation tests (with 999 randomization) were used to assess the overall significance of the analysis, the significance of each constrained axis, as well as the significance of each term using the function ‘anova.cca’. We used the ‘varpart’ function of ‘Vegan’ to partition the variation in earthworm community data with respect to the main explanatory variables highlighted by the RDA.

In order to highlight potential indirect effects of environmental variables on earthworm community structure, we performed path analyses using the ‘psem’ function of the ‘piecewiseSEM’ package [56]. To fix the theoretical model required for these analyses, we first explored all possible correlations between both community (*Str.1* and *Str.2*) and environmental (*Clim.1, Soil.1, Soil.2, Veg1.1, Veg1.2, Veg2.1, Veg2.2*) synthetic variables. We thus built a theoretical complete model figuring all the potential interactions between community and explanatory variables, and ran the path analysis using generalised linear models (glm) to build the list of structural equations.

## 3. Results

### 3.1. Species richness and community composition

We found a total of six genera and 11 species of earthworms across the whole altitudinal gradient (Fig. 1, Table A.2). While most mean species occurrences were below the treeline (2000-2100 m), seven species were distributed in both forest ecosystems and alpine meadows. The species *Dendrodrilus subrubicundus* (Eisen, 1874) was the most alticolous in our survey, with a mean occurrence just above 2200m.

**Figure 1.**
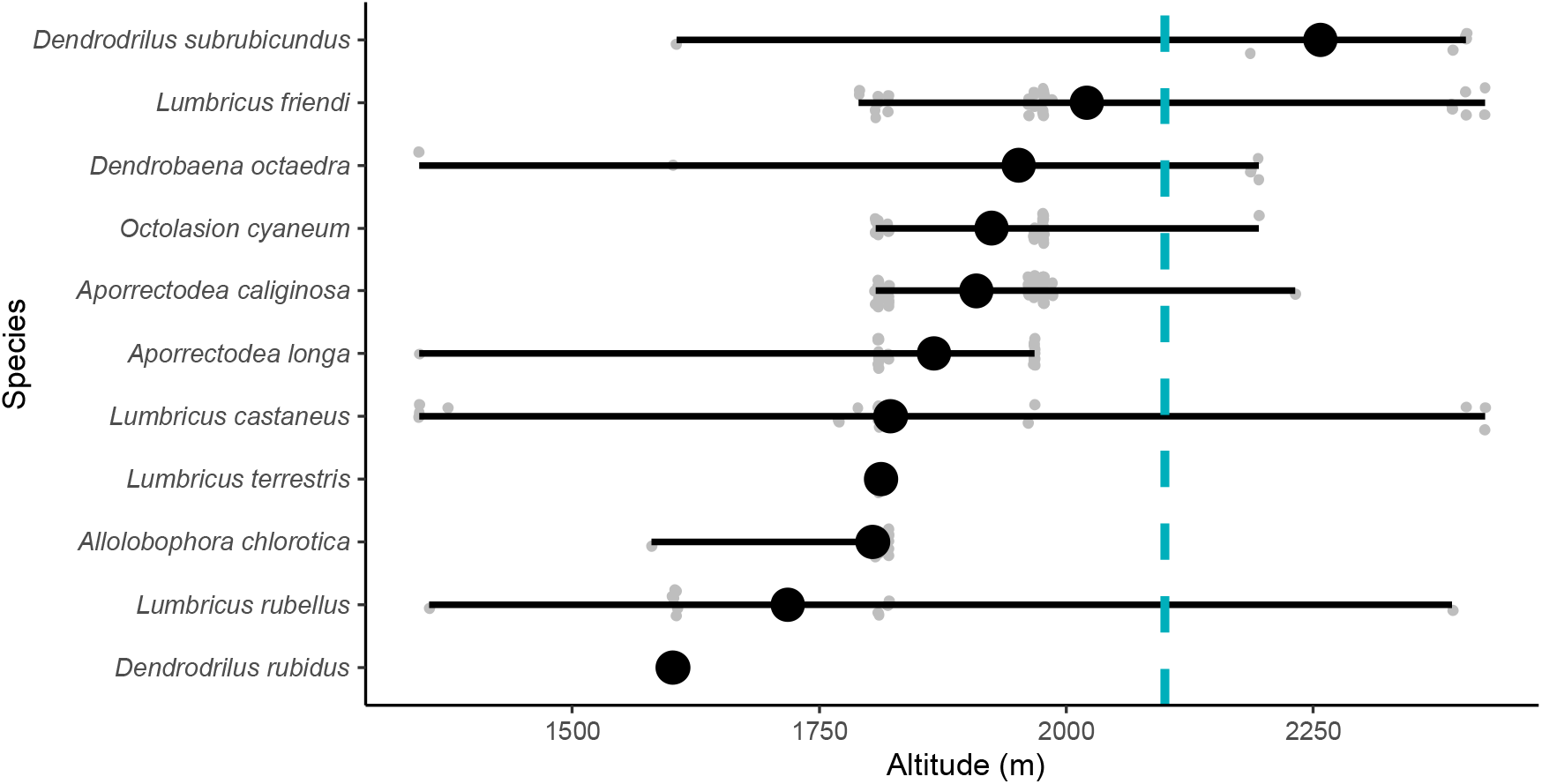
List of the earthworm species found in Belledonne Massif (France) and their altitudinal ranges. Bars indicate the minimal and maximal altitudes where each species was collected, large black dots represent the mean altitude, and small grey dots the individual occurrences with a jitter-randomized position allowing them to be viewed without overlap. The vertical blue line represents the altitudinal position of the treeline.

The altitudinal variation of earthworm species density (number of species observed per sample) and cumulated richness at the scale of each altitudinal stage are shown in Fig. 2. Species density ranged from 0 to 7 species per sample (mean ± sd: 1.4 ± 1.7; Fig. 2A), and cumulated species richness ranged from 5 to 9 species per altitudinal stage (Fig. 2B). Species diversity generally peaked at the ecotone just below the treeline between 1800 and 2200 m of altitude. This trend was significant while considering species density, but it was partly masked when considering cumulated richness by the high inter-plot variability in species composition within each altitudinal stage. The Chao asymptotic richness showed a similar pattern (Table 2).

**Figure 2.**
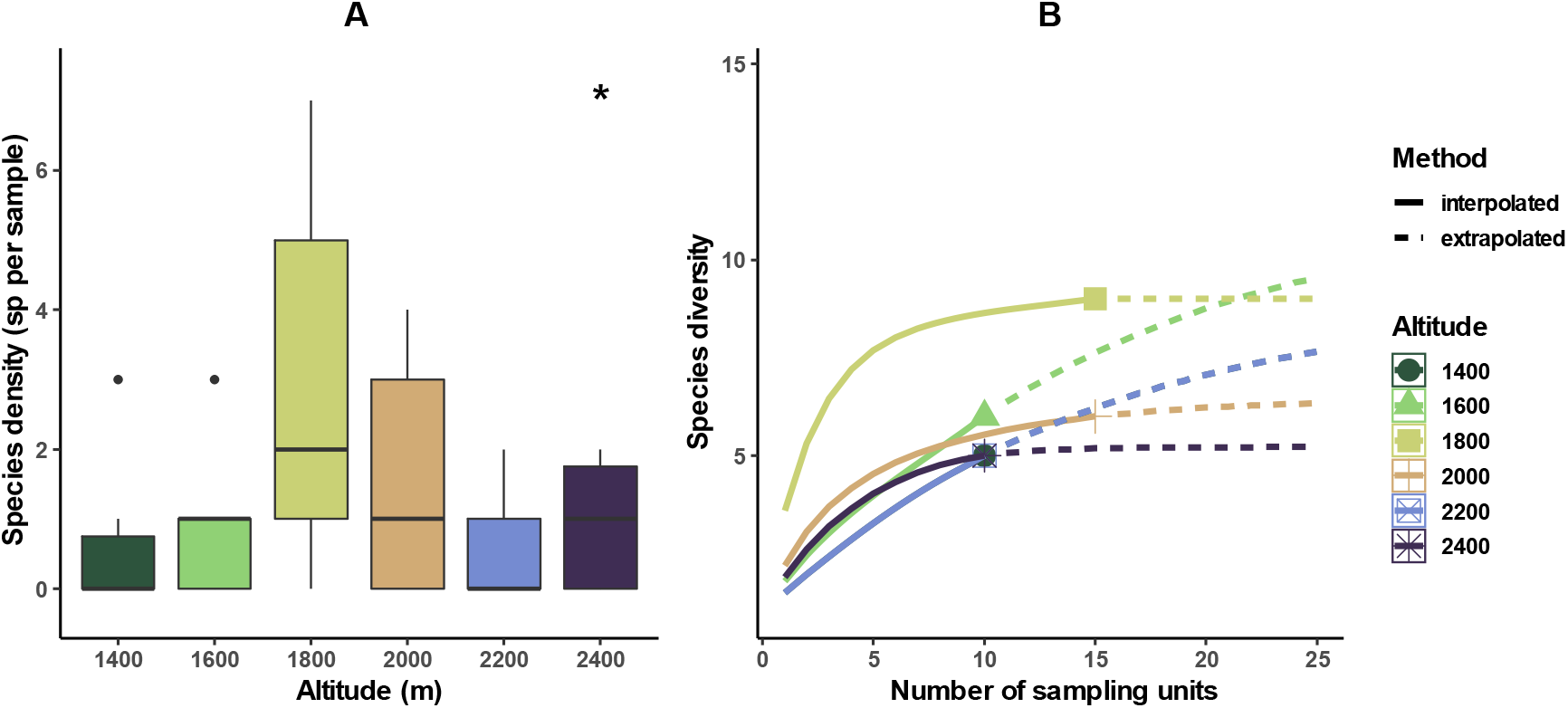
Variation of earthworm species diversity along the altitudinal gradient: A) Box-plots showing the variation of the average species density per altitudinal stage; B) Sample-based rarefaction and extrapolation curves of species diversity plotted for each altitudinal stage. In A) black dots are outliers; significance codes for Kruskal-Wallis rank sum test: 0 ‘***’ 0.001 ‘**’ 0.01 ‘*’ 0.05 ‘NS’. In B) solid lines represent rarefaction (interpolated) curves, whereas dashed lines represent extrapolated curves; shaded areas are 95% confidence intervals based on a bootstrap with 200 replications.

**Table 2.** Cumulated observed (S.obs) and estimated (S.Chao) earthworm species richness along the altitudinal gradient. S.Chao.se = standard error of the Chao’s asymptotic estimator.

| Altitudinal level (m) | S.obs | S.Chao | S.Chao.se |
| --- | --- | --- | --- |
| 1400 | 4 | 6.57 | 3.75 |
| 1600 | 5 | 10.46 | 6.39 |
| 1800 | 8 | 8.00 | 0.10 |
| 2000 | 5 | 5.00 | 0.34 |
| 2200 | 4 | 6.57 | 3.75 |
| 2400 | 4 | 4.00 | 0.51 |

The Kruskal-Wallis test did not highlight any significant effect of vegetation composition (structuring species) on earthworm species density (result not shown), while on the contrary the effect of altitude was significant (p-value < 0.01). The pairwise Wilcoxon rank sum tests show that this effect was essentially driven by the specific diversity at 1800 m, which differed significantly from that of all the other altitudes.

We found a significant pattern of variation of epigeic and endogeic scores along the altitudinal gradient, while no significant trend was observed for anecics (Fig. 3). The pairwise Wilcoxon rank sum tests highlight that this was mainly due to communities at 1800 and 2200 m of altitude, which had a significantly lower epigeic score and a significantly higher endogeic score compared with those at other altitudinal levels. No effect of vegetation was detected on ecological category scores (p-value > 0.05; result not shown).

**Figure 3.**
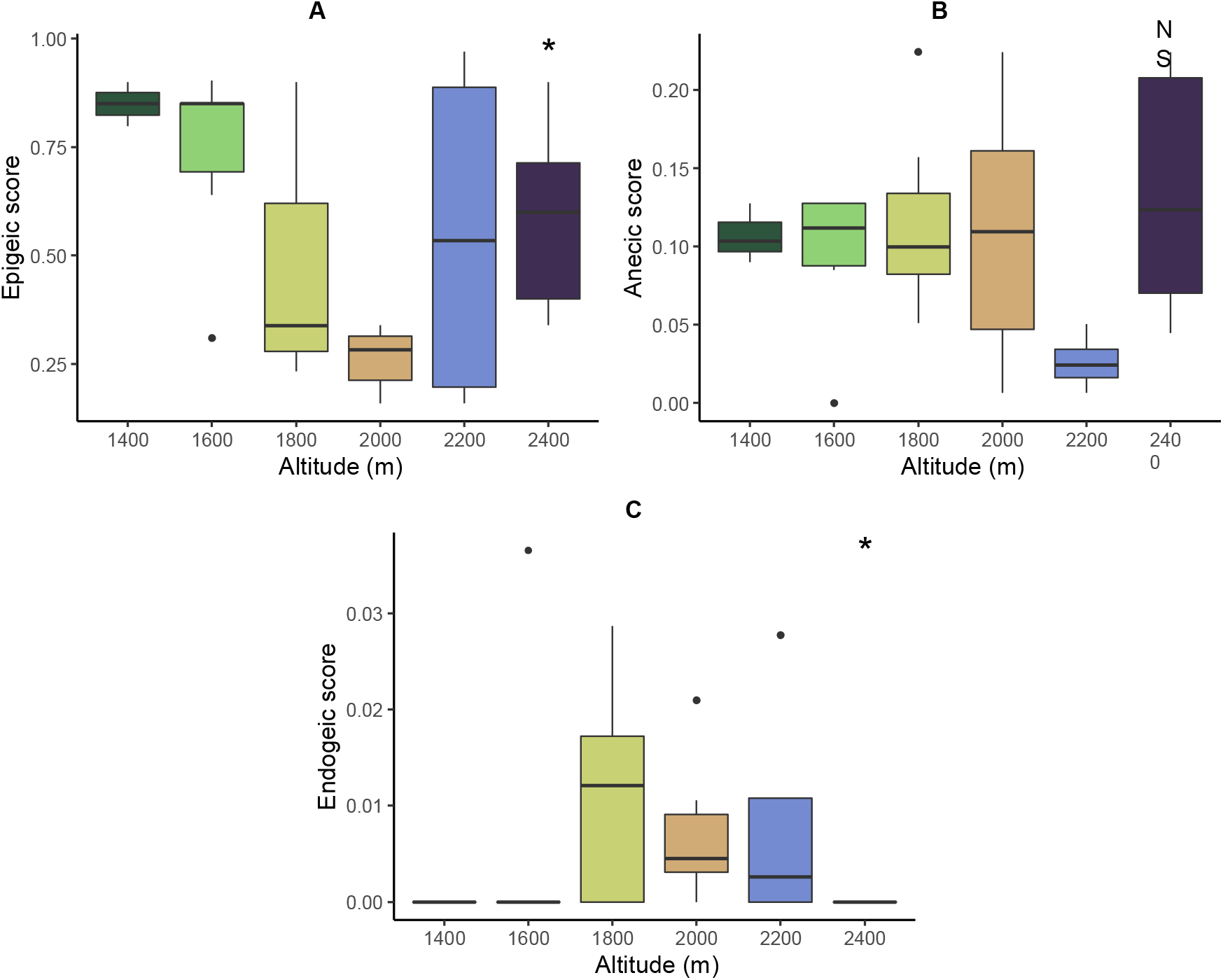
Variation of the community weighted mean of ecological categories scores along the altitudinal gradient: A) Epigeic score, B) Anecic score, C) Endogeic score. Significance codes for Kruskal-Wallis rank sum test: 0 ‘***’ 0.001 ‘**’ 0.01 ‘*’ 0.05 ‘NS’.

### 3.2. Density, biomass and body mass

Community density ranged from 0 to 416 individuals per m^2^ (mean ± sd: 51.4 ± 86.0), and biomass ranged from 0 to 135 g per m^2^ (mean ± sd: 12.7 ± 25.2). Both density and biomass showed a similar pattern of variation along the altitudinal gradient, with a peak reached at the transition zone (Fig. 4A-B). The Kruskal-Wallis tests showed no significant effect of vegetation (p-value > 0.05), while altitude significantly explained the variability in both biomass and density (p-value < 0.01 and 0.05, respectively).

**Figure 4.**
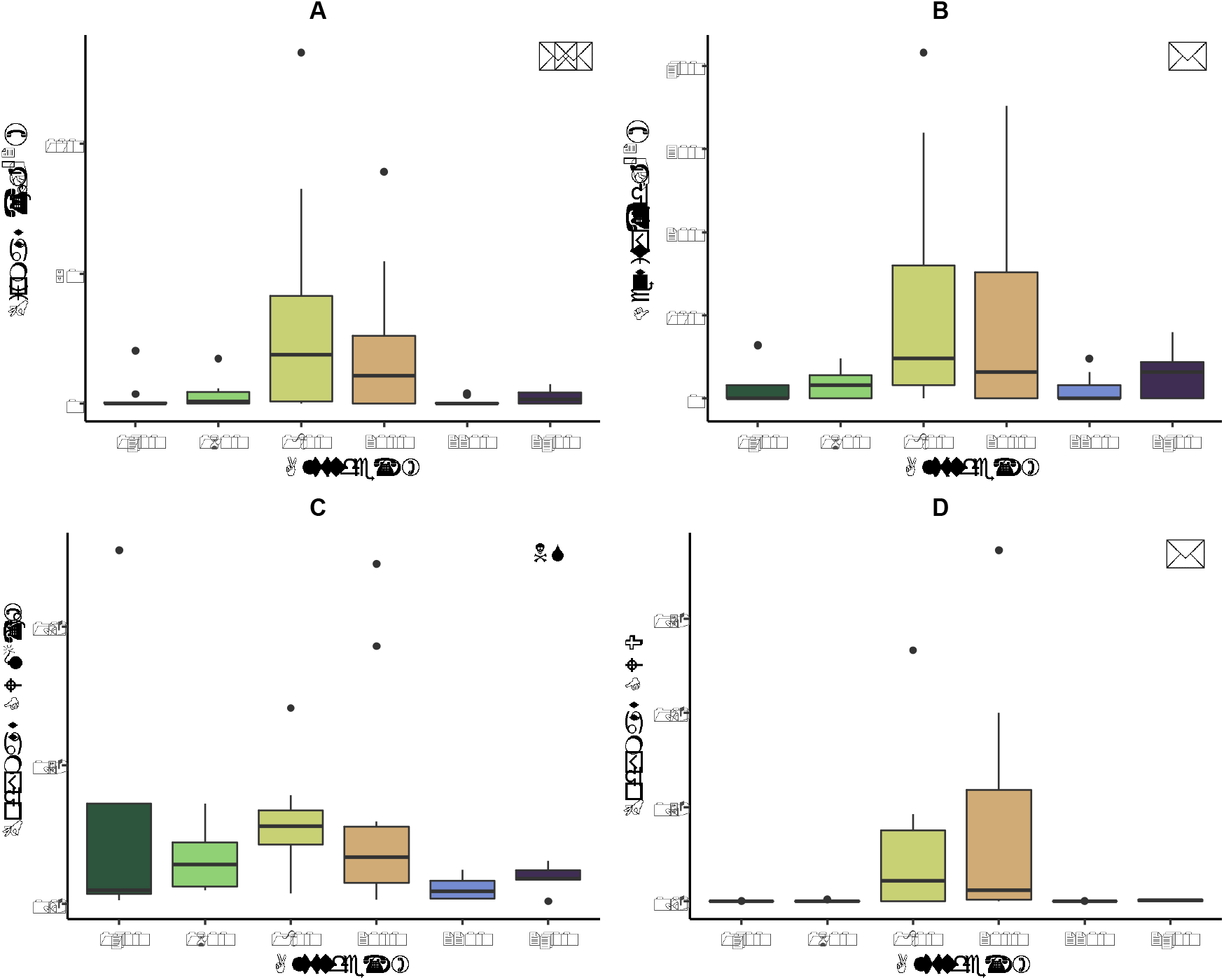
Variation of biomass (A), density (B), body mass community weighed mean (C) and community weighted variance (D) along the altitudinal gradient. Significance codes for Kruskal-Wallis rank sum test: 0 ‘***’ 0.001 ‘**’ 0.01 ‘*’ 0.05 ‘NS’.

No significant variation in body mass CWM was observed along the altitudinal gradient (Fig. 4C.), and no effect of vegetation type was also found (p-value > 0.05; result not shown). On the contrary, body mass CWV varied along the gradient in a significant way and according to a pattern comparable to that observed for biomass and density, with values peaking at 1800 and 2000m of altitude (Fig. 4D).

### 3.3. Environmental drivers of community structure

The redundancy analysis was significant after the permutation test was applied (p-value < 0.001). The first two axes explained 58.4% and 19.8% of the covariation between community and environmental tables (Fig. 5A). The proportion of constrained inertia was of 43.5% (Fig. 5D). Only the first axis appeared to be significant after the permutation test was applied (p-value < 0.001; Fig. 5E).

**Figure 5.**
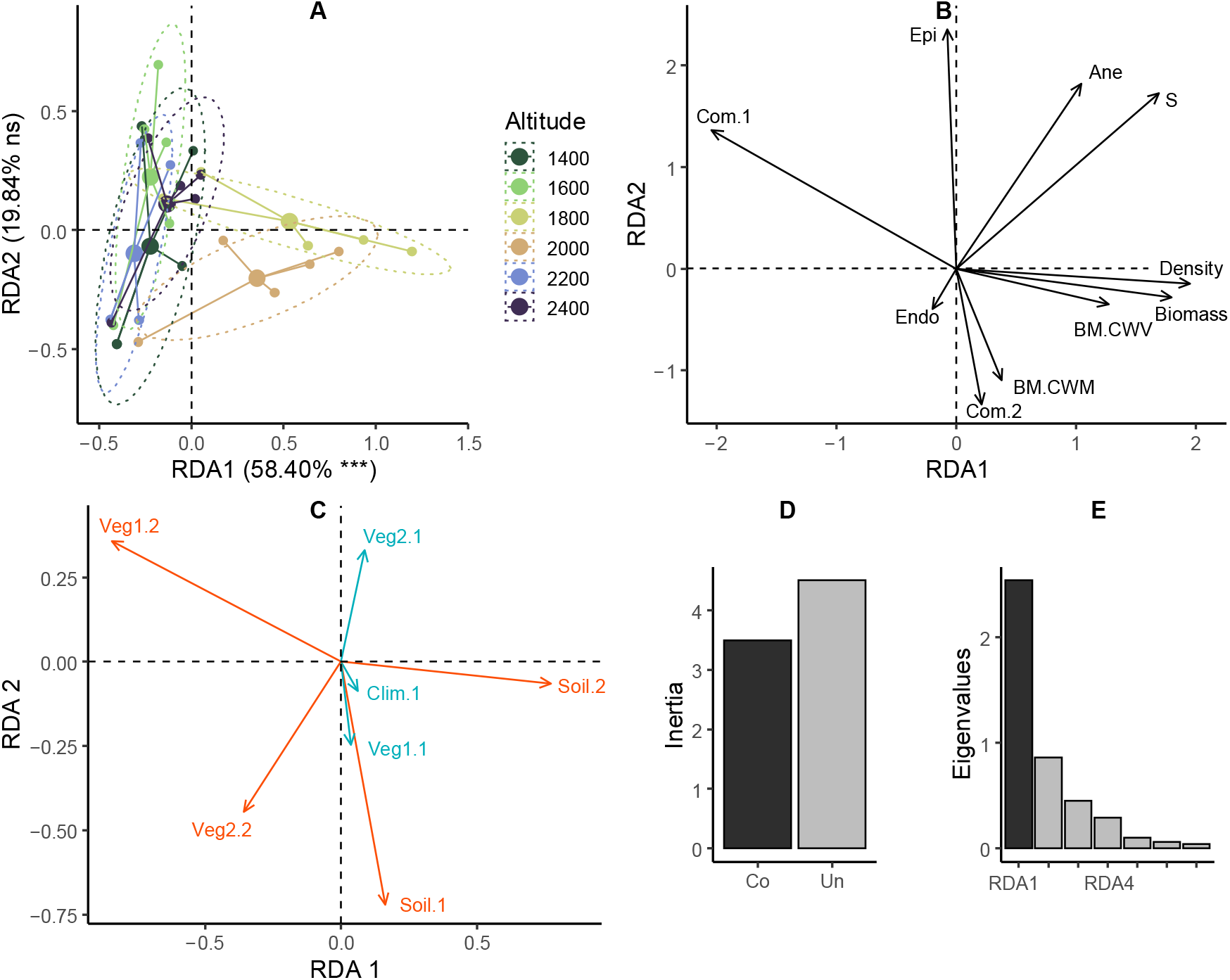
Redundancy analysis (RDA) highlighting the relative importance of environmental predictors in explaining altitudinal changes in earthworm community structure and composition: A) ordination of sampling plots, groups by altitudinal levels, on the factorial plan defined by the first two axes of the RDA (i.e. RDA1 and RDA2); B) projection of community metrics on the first factorial plan; C) projection of synthetic environmental predictors on the first factorial plan; D) relative proportion of constrained (Co) and unconstrained (Un) inertia; E) eigenvalue diagram. In A) significance codes for the RDA axes after permutation test: 0 ‘***’ 0.001 ‘**’ 0.01 ‘*’ 0.05 ‘ns’. In B) S= observed species richness; BM= body mass; CWM= community weighted mean; CWV= community weighed variance; Endo, Epi & Ane= CWM of endogeic, epigeic and anecic scores; Com.1 & Com.2= plot scores on the first two axes of the PCA of earthworm community composition (see Table 1). In C) synthetic environmental predictors correspond to the plot scores on the first two axes of the PCAs of climatic variables (Clim.1 & Clim.2), soil properties (Soil.1 & Soil.2), vegetation structure (Veg1.1 & Veg1.2) and vegetation composition (Veg2.1 & Veg2.2) (see Table 1); red arrows indicate significant predictor variables after permutational ANOVA (p-value < 0.05).

The first RDA axis highlighted the ecotone effect in the structuring of communities. Indeed, plots located in the transition zone just below the treeline (i.e. 1800 and 2000m altitudinal levels) had positive scores on this axis, and were clearly opposed to all other altitudinal levels (Fig. 5A). The projection of the community variables highlighted that earthworm communities in these plots were characterized by higher density, biomass and species richness, more variable body mass among co-existing species, and community composition dominated by species displaying negative scores on the community component *Com.1* (essentially *Lumbricus terrestris* L., 1758 and *Octolasion cyaneum* (Savigny, 1826); Fig. 5B, Fig. A.4B). The projection of environmental variables on RDA1 showed this ecotone effect was mainly driven by soils richer in nitrates and with a higher pH (positive projection of *Soil.2* on RDA1), and by a greater representation of the herbaceous stratum (grasses and forbs) in the vegetation structure (negative projection of *Veg1.2* on RDA1; Fig. 5C, Fig. A.3F).

Even though axis 2 of the RDA was not significant after the permutation test (p-value = 0.18), we decided to keep it in the interpretation because it highlighted an effect of the altitudinal gradient that could be partially hidden by the dominant ecotone effect. Plots located below 1800m had positive averaged scores on RDA2, while plots above 2000m displayed negative scores (Fig. 5A). The projection of community variable on this axis showed that lower elevation levels hosted communities with a higher average epigeic score, mainly because of the presence of *Lumbricus rubellus* (Hoffmeister, 1843), *Dendrobaena octaedra* (Savigny, 1826) and *Dendrodilus rubidus* (Savigny, 1826), and lower body mass CWM (Figs. 5B, 5C, Fig. A.4B). This effect was mainly related to an altitudinal gradient in soil properties (*i.e. Soil.1*; Fig. 5C), i.e. an increase in sand content and bulk density with altitude, accompanied by a decrease in organic matter and nutrient content (Fig. A.4D). Vegetation structure was also linked, albeit less clearly, to RDA2, mainly because of a gradual replacement of *J. communis* by other shrubs with increasing altitude (*i.e. Veg2.2*; Fig. 5C, Fig. A.4H).

The variance partitioning analysis highlights that a large proportion of the variance in earthworm community structure is not explained by the monitored environmental drivers (residuals = 73.3%; Fig. A.5). Among the explained fraction of the variance, a significant proportion (i.e. 16.4% of the total variance) was explained by the interaction between the variables *Soil.2* and *Veg1.2*. Then, the main environmental predictors, in decreasing order of importance, were *Veg1.2, Soil.1, Veg2.2* and *Soil.2* (Fig. A. 5).

The path analyses highlight the main direct and indirect environmental controls on earthworm communities. The ecotone effect on community structure (*Str.1*), which is described by the first axes of both the RDA (Fig. 5B) and the PCA of community structure (Figs. A.4C-D), is firstly explained by an indirect effect of soil nitrate concentration and pH (*Soil.2*) mediated by the importance of the herbaceous stratum in vegetation structure (*Veg1.2*), and secondly by a direct effect of the composition of the shrub stratum in the vegetation (*Veg2.2*) (Fig. 6A). The altitudinal effect on community structure (*Str.2*), which is described by the second axes of both the RDA (Fig. 5B) and the PCA of community structure (Figs. A.4C-D), is explained by an effect of soil properties (i.e. altitudinal variations in sand content, bulk density, organic matter and nutrient contents; *Soil.1*) partially determined by climate (*Clim.1*), and by a direct effect of shrub stratum composition (*Veg2.2*) (Fig. 6).

**Figure 6.**
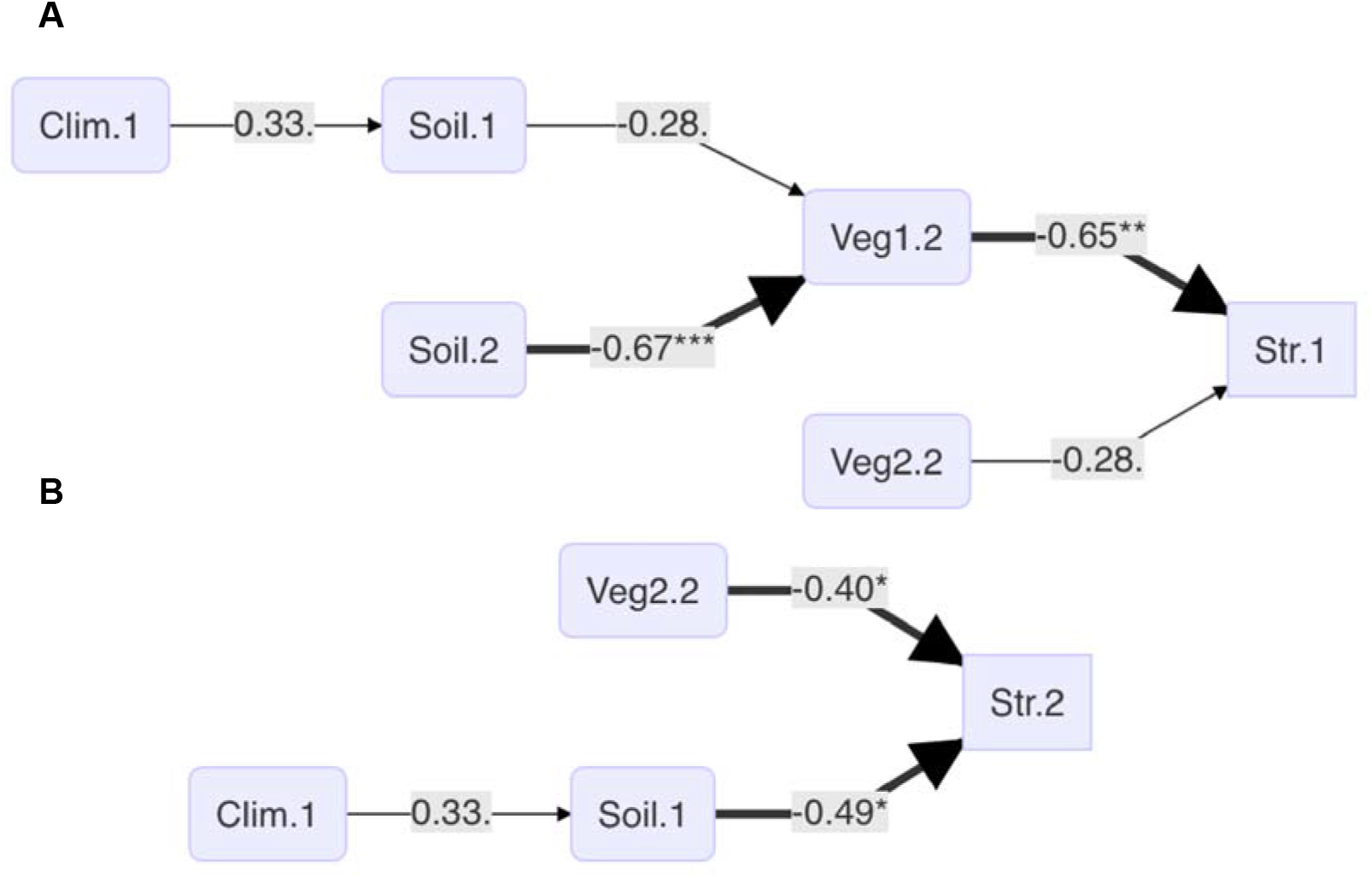
Path analyses showing the direct and indirect effects of environmental predictors on earthworm community structure. Str.1 and Str.2 correspond to the plot scores on the first two axes of the PCA of community structure, respectively (see Fig. A.4D). Significance codes for thick arrows: 0 ‘***’ 0.001 ‘**’ 0.01 ‘*’ 0.05; thin arrows indicate marginally significant correlations (p-value < 0.1). See Table 1 for interpretation of the synthetic environmental and community synthetic descriptors.

## 4. Discussion

The mean values of density and biomass (51 individuals per m^−2^, and 12 g.m^−2^, respectively), and species richness (1.4 species per sample, and 5 to 7 species per altitudinal stage) that we found in our study are within the range of what is generally reported for comparable temperate ecosystems. Earthworm communities in montane or boreal coniferous forests, for example, have abundances ranging from 0 to about 100 individuals, m^−2^, biomasses ranging from 0 to 25 g. m^−2^, and are generally composed of fewer than 5 species [57–60]. Similarly, in mountain pastures, earthworm density can vary from 60 to 200 individuals per m^−2^, biomass from 15 to 30 g. m^−2^, and communities generally comprise 2 to 4 species [60,61]. These figures are on average lower than those generally given for lowland forest and grassland ecosystems [62–65]. The fact that earthworm communities are less abundant and less diverse in the mountains than in the lowlands can be explained by the unfavourable climatic conditions, the vegetation dominated by species producing high C:N litter (especially conifers), as well as by the presence of shallow soils in the higher altitudinal levels of the altitudinal gradient.

Firstly, our study does not show a monotonic decrease in species richness along the altitudinal gradient. Few studies have looked at the variation in species diversity of earthworm assemblages along altitudinal gradients. Some have described a monotonic decrease in species number with altitude, for example in France [37,66] or in Serbia [67]. It should be noted, however, that these previous studies considered a complete altitudinal gradient (i.e. from sea level to alpine levels), which is not the case in our study. In another study in Puerto Rico, Gonzales et al. [68] observed, on the contrary, an increase in the number of earthworm species along an altitudinal gradient from 0 to 1000 m, which the authors interpreted as the result of the combination of several soil and climatic factors. Finally, Fontana et al. [20] described a pattern similar to ours along a gradient from 1000 to 2500 m in northern Italy, with a peak in species diversity observed between 1800 and 2200 m altitude. Our results therefore confirm the latter study and largely invalidate the hypothesis of a monotonic effect of altitude and climate. The absence of a monotonic decrease in the number of species with altitude can be explained in our case by the composition of the regional species pool, which is essentially dominated by species with a wide distribution in Northern Europe. Indeed, after the last Würm glaciation, the areas left free by the retreat of glaciers were recolonised by species with a high dispersal capacity and ecological plasticity [37]. This mechanism occurred simultaneously along a latitudinal gradient towards northern Europe, allowing the establishment of earthworm communities up to the present-day boreal forests, and in mountain areas up to the alpine levels. We therefore do not observe the increase in endemism classically observed with altitude for other taxa [69], nor do we observe a steady decrease in species richness along our altitudinal gradient.

Our study highlights the importance of the ecotone associated with the treeline as the primary factor responsible for the structuring of earthworm communities. At the level of this ecotone, earthworm communities are more abundant and diverse, and show a higher CWV of body mass, notably due to the presence of large species such as *L. terrestris* and *O. cyaneum* co-occurring with smaller body-sized species such as *Aporrectodea caliginosa* (Savigny, 1826) and *Lumbricus castaneus* (Savigny, 1826). The observation of a peak in species richness in the forest-grassland transition zone at around 1800–2000 m altitude is at first sight suggestive of the mid-domain effect, which predicts a higher species diversity in this zone due to the overlapping altitudinal distributions of certain species [70,71]. However, this interpretation does not seem convincing in the context of our study, given the already mentioned broad ecological spectrum that characterizes the species composing the regional pool. Indeed, it is unlikely that the species present along our gradient would show a preference for a particular altitudinal zone, and the peak in diversity observed at the ecotone level must in our view be attributed to ecological factors rather than to a geometric overlap of species distributions.

According to our results, community structure at the ecotone is associated with high nitrate and soil pH, but this effect of soil properties is indirect and largely mediated by vegetation structure. In particular, the presence of herbaceous cover and shrub layer appears to be the main determinant of earthworm communities. This result contrasts with the findings of other works that emphasize soil properties (including pH, C:N, texture and depth) as the dominant factor in explaining the distribution of earthworm abundance, biomass or diversity [58,59,63]. However, Phillips et al. [72] demonstrated that the effect of soil properties on earthworm communities may be scale-dependent, and that it may be hidden by other factors such as habitat cover or annual rainfall depending on the study context. In our study, the deliberate inclusion of the ecotone in the sampling design, and the fact that this area clearly contrasted environmentally with the other parts of the altitudinal gradient, probably contributed to emphasizing vegetation structure rather than soil properties, whose direct effect is relegated to axis 2 of the RDA. Moreover, the use of path analyses is not common in studies on the driving factors of earthworm communities, whereas in our study they allowed to highlight this indirect effect. The generalization of this type of approach would probably allow a better understanding of the complex pathways of environmental controls on earthworm communities.

Several studies have highlighted that transitional ecosystems represent favourable habitats for earthworms. Bernier and Ponge [73] and Grossi & Brun [74] had shown that along mountain succession gradients, transitional phases between pastures and forests host abundant and diversified earthworm communities. In lowland environments, other studies have described the same type of pattern in the early stages of post-pastoral or post-cultural successions [26,27]. This pattern can firstly be explained by a greater diversity of micro-habitats at the level of the ecotone, compared to that of the ecosystems of which it marks the transition [1,4]. For earthworm communities, the ecotone provides alternating patches of herbaceous and woody vegetation, which influence soil microclimate and in which litter accumulations vary in thickness and quality depending on the identity of the plant species (i.e. coniferous versus deciduous). This environmental variability allows species with different ecological requirements to co-exist in the same communities. This point, which is consistent with our hypothesis of habitat complexity, is further supported in our results by the observed peak in body-mass CWV at the ecotone, which suggests that earthworm communities are morphologically and therefore functionally more diverse in this area of the gradient.

Furthermore, the effect of the ecotone can be linked to the transition from a vegetation dominated exclusively by conifers to a mixed vegetation, i.e. with a significant proportion of deciduous species. A reduced proportion of conifers in the vegetation can have an impact on earthworm communities through an improvement of the quality of trophic resources. Indeed, it has been shown in several studies that mixed forests can host more abundant and diverse earthworm communities than coniferous forests can [58,75]. The openness of the canopy at the ecotone level also allows the establishment of dense herbaceous vegetation, which produces a litter layer rich in nutrients such as calcium and nitrogen [22,63,76]. Herbaceous species also participate, through both the quality of their organic matter input and their dense root mat, in the formation of a deep A horizon that creates suitable conditions for the activity of endogeous species [22,27,62]. This is supported in our results by the average score of the endogeic ecological category peaking at the ecotone. The general improvement of habitat suitability is further supported by the highest earthworm abundance and biomass observed at both 1800 and 2000 m of elevation. In general, our results show that, in addition to the environmental heterogeneity hypothesis, the structuring of earthworm communities in response to the tree-line can also be explained using the resource quality hypothesis.

Compared to the effect of the ecotone, altitude as such is of less importance in our results. However, earthworm communities at lower altitudes show a higher epigeic score and a lower CWM of body mass, even though we found no evidence that Bergmann’s law applies to our case study. This tendency for an altitudinal effect is mainly related to soil properties, i.e. an increase with altitude in sand content and bulk density, followed by a decrease in organic matter and nutrient content, and an increase in soil C:N. A smaller part of the variability in community structure is further explained by climate and vegetation composition. These results confirm the already observed trend towards low-abundant and epigeous-dominated earthworm communities in coniferous forests, as a consequence of low soil pH and high soil C:N in this type of forests [63,64,75]. Soil texture is another factor recognized for its importance in community structuring, especially in mountain environments. Salomé et al. [59] showed that epigeic species were more abundant in shallow, sandy soils, whereas anecics and endogeics were more abundant in deeper, finer-textured soils. Behind the dominant effect of the ecotone associated with the tree-line, altitudinal variability in soil properties, rather than the effect of altitude as such, can explain a moderate fraction of the variability in earthworm community structure. As already suggested earlier, it is possible that this hierarchy of driving factors is scale-dependent, and that soil factors might be of primary importance if the sampling design had been designed to explain community structuring within each altitudinal stage rather than along the whole gradient.

Overall, our results largely invalidate the hypothesis of a monotonic effect of climate, but fail to separate the potential effect of environmental heterogeneity and resource quality, both of which can be invoked to explain the observed community pattern at the ecotone level. In the context of global change, an upward shift of the treeline is expected as a result of climate warming [77,78]. In contrast to what is predicted for other taxa [69,79], the prospect of this shift does not appear to represent an *a priori* threat to the diversity of earthworm species, which includes no montane endemics at our study site. However, the functional implications of these changes could be considerable. In particular, a shift of the ecotone to higher altitudinal levels would inevitably lead to a reduction in its surface area, and consequently in the size of the associated earthworm activity hotspot. In the future, it will be necessary to examine the consequences that this could have on ecosystem functioning at the scale of such mountain landscapes.

## 5. Acknowledgments

The authors acknowledge the Mairie de Chamrousse, Isère, France, for access to field sites, and all the participants to the ECOPICS project for their contribution to logistics during field sampling. The study was founded by the Agence Nationale de la Recherche from France and the Consejo Nacional de Ciencia y Tecnología from Mexico (ECOPICS project, ANR-16-CE03-0009 and CONACYT-2 73659).

