## Appendix for "Environmental drivers of earthworm communities along an altitudinal gradient in the French Alps"

***Appendices***

***Appendix 1 – Study sites***

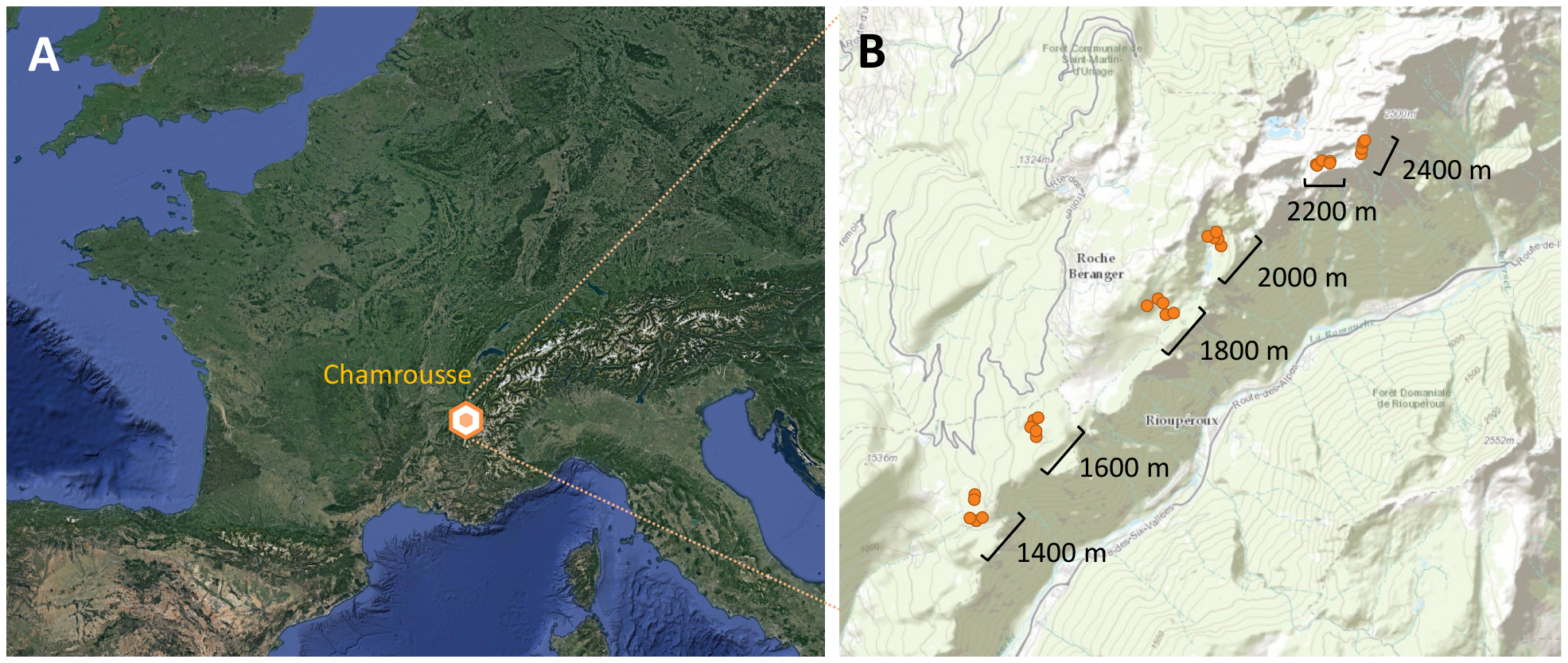
***Figure A.1.*** Map of the study sites situated along the altitudinal gradient in the Belledonne massif (France): A) Localisation of Chamrousse in France; B) Localisation of the sampling plots in the six altitudinal levels.

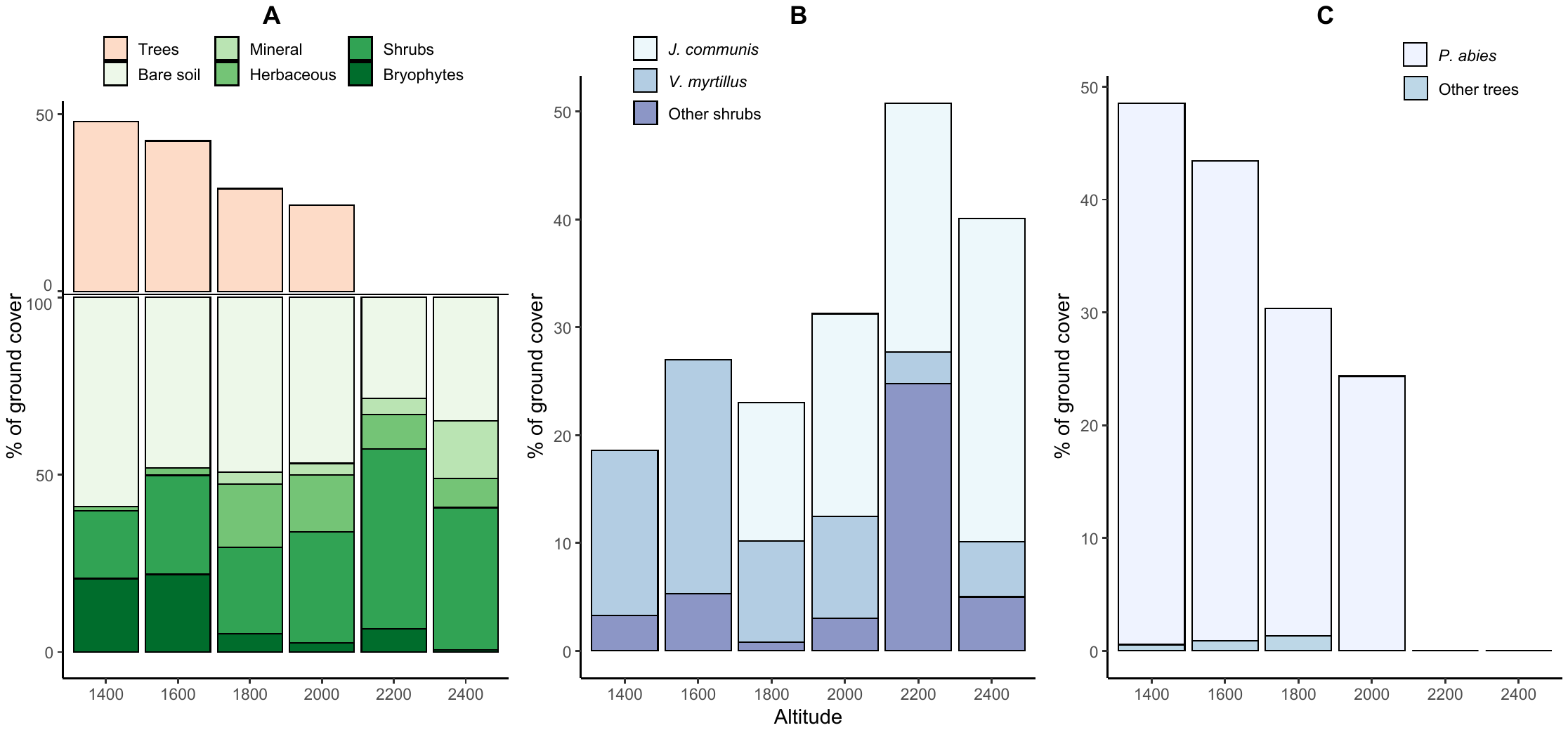

***Figure A.2.*** Vegetation characteristics along the altitudinal gradient: A) relative cover of different vegetation strata; B) relative cover of the two shrub target species (*Juniperus communis* and *Vaccinium myrtillus*) within the shrub layer; C) relative cover of *Picea abies* within the tree cover (modified after [1]).

***Appendix 2 – Environmental variables***

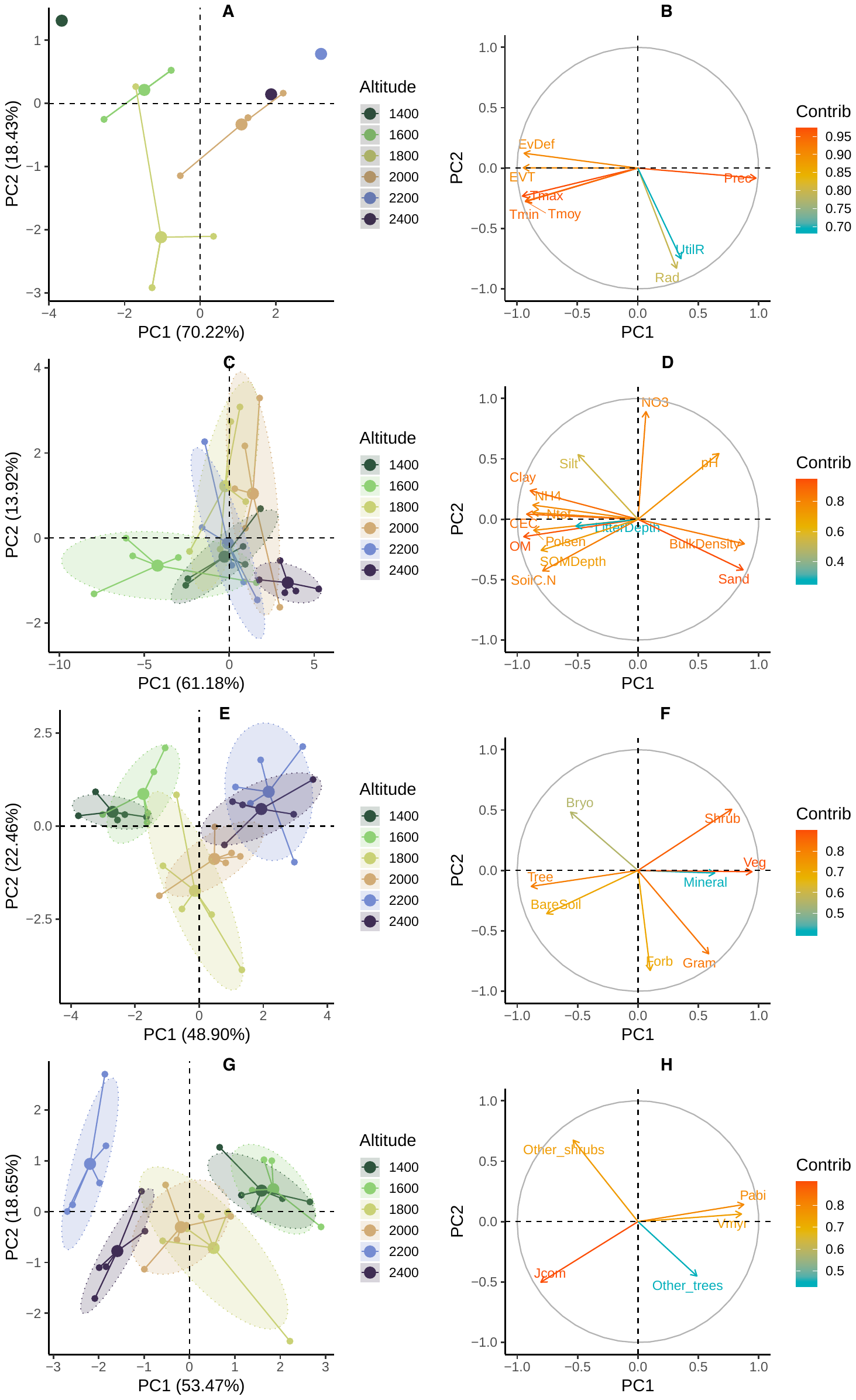

***Figure A.3*** (previous page)***.*** Principal component analyses (PCA) of environmental tables: A-B) climate variables, C-D) soil properties, E-F) vegetation structure, G-H) vegetation composition. For each individual PCA, the left figure represents the ordination of sampling plots, grouped by altitudinal levels, on the factorial plan defined by the first two axes, and the right figure represents the projection of the elementary variables on the same factorial plan (see Table A.1 for the meaning of the elementary variable codes).

| **Codes** | **Variable** | **Unit** | **Codes** | **Variable** | **Unit** |
| --- | --- | --- | --- | --- | --- |
| Climate |  |  | Vegetation structure | | |
| *Tmax* | Mean annual maximal temperature | °C | *Mineral* | Percentage of Mineral Cover | % |
| *Tmoy* | Mean annual temparature | °C | *Bryo* | Bryophyte cover | % |
| *Tmin* | Mean annual minimal temperature | °C | *Veg* | Vegetation cover | % |
| *Rad* | Mean radiation | W | *BareSoil* | Bare soil cover | % |
| *EVT* | Potential evapotranspiration | mm | *Forb* | Forb cover | % |
| *Prec* | Mean annual precipitations | mm | *Gram* | Graminoid cover | % |
| *EvDef* | Evaporation deficit | mm | *Shrub* | Shrub cover | % |
| *UtilR* | Usefull water reserve | mm | *Tree* | Tree cover | % |
| Soil properties | |  | Vegetation composition | | |
| *Clay* | Clay proportion | g/kg | *Vmyr* | *Vaccinium myrtillus* cover | % |
| *SiltFine* | Fine silt proportion | g/kg | *Jcom* | *Juniperus communis* cover | % |
| *SiltCoarse* | Coarse silt proportion | g/kg | *Pabi* | *Picea abies* cover | % |
| *SandFine* | Fine sand proportion | g/kg | *Other_trees* | Cover by other tree species than *P. abies* | % |
| *SandCoarse* | Coarse sand proportion | g/kg | *Other_shrubs* | Cover by other shrub species than *V. myrtillus* and *J. communis* | % |
| *SOC* | Soil organic carbon | g/kg |  |  |  |
| *Ntot* | Total nitrogen | g/kg |  |  |  |
| *OM* | Organic matter | g/kg |  |  |  |
| *SoilC.N* | C:N ratio |  |  |  |  |
| *pH* | pH |  |  |  |  |
| *Polsen* | Phosphorus Olsen concentration | g/kg |  |  |  |
| *CEC* | Cationic exchange capacity | cmol+/kg |  |  |  |
| *NH4* | Ammonium concentration | g/kg |  |  |  |
| *NO3* | Nitrate concentration | g/kg |  |  |  |
| *LitterDepth* | Litter layer depth | cm |  |  |  |
| *SOMDepth* | Organo-mineral layer depth | cm |  |  |  |
| *BulkDensity* | Soil bulk density | g/cm3 |  |  |  |

***Table A.1.*** List of the elementary environmental variables used to describe the climate, the soil properties and the vegetation structure and composition in the different sampling plots.

The first axis of the principal component analysis of climate variables (i.e. *Clim.1* and *Clim.2*) explained 70.2% % of the total variance (Fig. A.3A). We only retained *Clim.1* for interpretation, which organised the sampling plots from low elevation (negative scores) to high elevation (positive scores; Fig. A.3B). The projection of the elementary variables on this axis showed that along the altitudinal gradient temperature and potential evapotranspiration decreased while mean annual precipitations increased (Fig. A.3B).

The first two axes of the principal component analysis of soil properties (i.e. *Soil.1* and *Soil.2*) explained 61.2% and 13.9% of the total variance (Fig. A.3C). *Soil.1* mostly highlighted an opposition between the 1600m (negative scores) and the 2400m (positive scores) altitudinal levels (Fig. A3C). The projection of the soil variables on this axis showed that lower elevation plots were characterised by a silty texture and higher organic and nutrient contents, while high elevation soils were characterised by a sandy texture, and higher bulk density and pH (Fig. A.3D). *Soil.2* opposed soil located in the transition zone (1800 and 2000m; positive scores) to the other altitudinal levels (negative scores; Fig. A.3C) and was mostly explained by higher nitrate contents observed in the transition zone (Fig. A.3D).

The first two axes of the principal component analysis of vegetation structure (i.e. *Veg1.1* and *Veg1.2*) variables explained 48.9% and 22.5% of the total variance (Fig. A.3E). *Veg1.1* organised plots according to the altitudinal gradient from those located in the 1400m level (negative scores) to those at 2400m (positive scores; Fig. A.3E). The projection of the elementary variables showed that this axis corresponded to a decrease in tree and bare soil cover and an increase in shrubs and graminoid vegetation cover (Fig. A.3F). *Veg1.2* opposed soil located in the transition zone (1800 and 2000m; negative scores) to the other altitudinal levels (positive scores; Fig. A.3E) and was mostly explained by higher forb and graminoid cover in the transition zone (Fig. A.3F).

The first two axes of the principal component analysis of vegetation composition (i.e. *Veg2.1* and *Veg2.2*) explained 53.5% and 18.6% of the total variance (Fig. A.3G). *Veg2.1* organised plots according to the altitudinal gradient from those located in the 1400m level (positive scores) to those at 2400m (negative scores; Fig. A.3G). The projection of the elementary variables showed that this axis corresponded to a decrease in *Picea abies* and *Vaccinium myrtillus*, paralleling an increase in *Juniperus communis*, from low to high elevation plots (Fig. A.3H). *Veg2.2* opposed plots located in 1400, 1600 and 2200m altitudinal levels (positive scores) to the other plots (negative scores; Fig. A.3G) and was mostly explained by higher ‘other shrubs’ cover in the former plots by opposition to *J. communis* cover (Fig. A.3H).

***Appendix 4 – Earthworm community structure***

| **Species Name** | **Code** | **Epigeic score** | **Anecic score** | **Endogeic score** | **Ecological category** |
| --- | --- | --- | --- | --- | --- |
| *Allolobophora chlorotica* | Achl | 0.31 | 0.31 | 0.38 | Intermediate |
| *Aporrectodea calliginosa* | Acal | 0.16 | 0.04 | 0.80 | Endogeic |
| *Aporrectodea longa* | Alon | 0.32 | 0.68 | 0.00 | Epi-anecic |
| *Dendrobaena octaedra* | Doct | 0.97 | 0.03 | 0.00 | Epigeic |
| *Dendrodrilus subrubicundus* | Dsub | 0.64 | 0.00 | 0.36 | Epi-endogeic |
| *Dendrodrilus rubidus* | Drub | 0.89 | 0.11 | 0.00 | Epigeic |
| *Lumbricus castaneus* | Lcas | 0.90 | 0.10 | 0.00 | Epigeic |
| *Lumbricus friendi* | Lfri | 0.34 | 0.66 | 0.00 | Epi-anecic |
| *Lumbricus rubellus* | Lrub | 0.85 | 0.15 | 0.00 | Epigeic |
| *Lumbricus terrestris* | Lter | 0.34 | 0.66 | 0.00 | Epi-anecic |
| *Octolasium cyaneum* | Ocya | 0.21 | 0.24 | 0.55 | Intermediate |

***Table A.2.*** List and ecological characteristics of the earthworm species observed in the study. Affinity scores with the three ecological categories of Bouché [2], and synthetic ecological category are defined according to Bottinelli et al. [3].

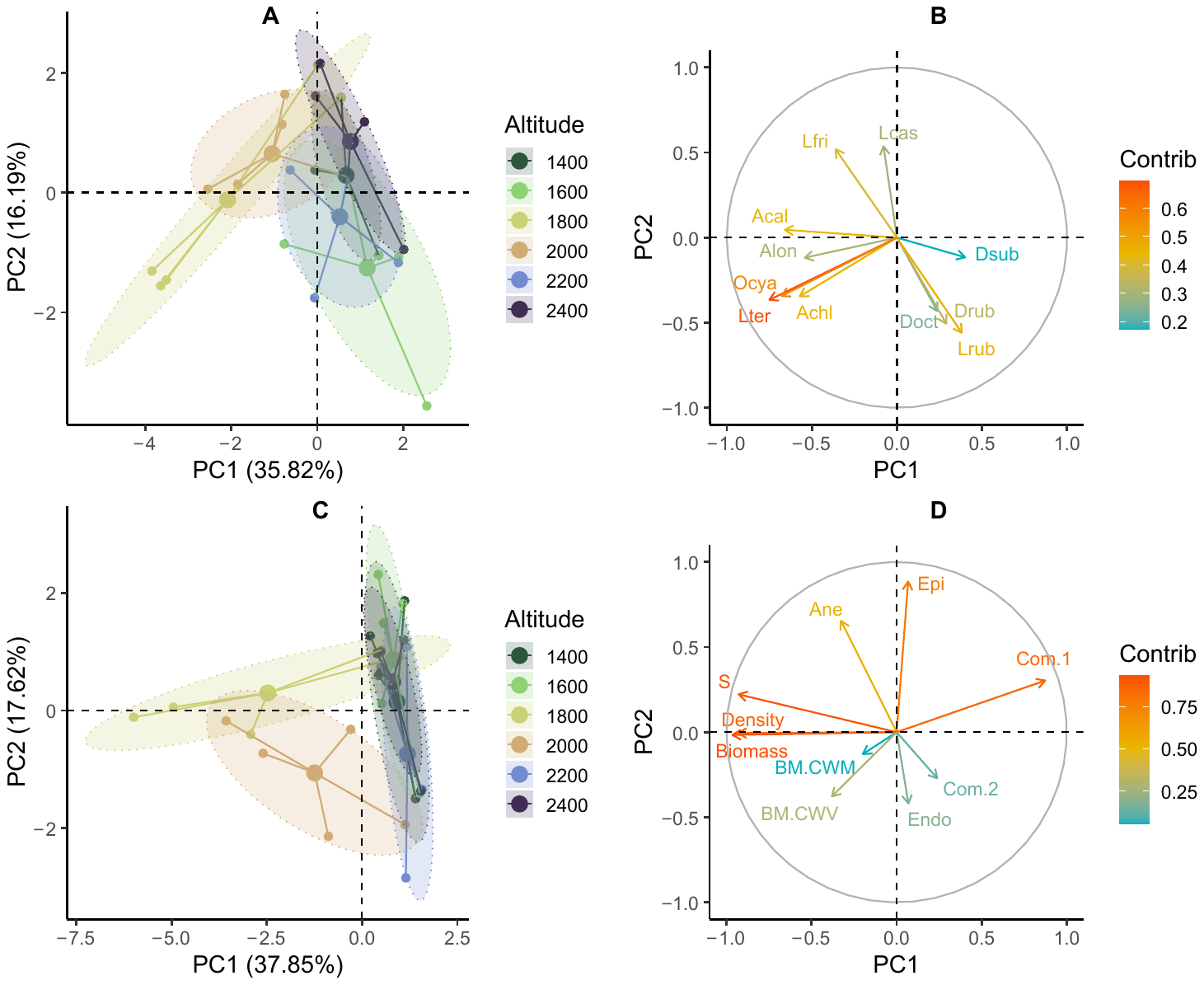

***Figure A4.*** Principal component analyses (PCA) of earthworm community composition and structure: A) ordination of sampling plots, grouped by altitudinal levels, on the factorial plan defined by the first two axes of the PCA of community composition (i.e. *Com.1* and *Com.2*); B) projection of species variables on the same factorial plan; C) ordination of sampling plots, groups by altitudinal levels, on the factorial plan defined by the first two axes of the PCA of community structure (i.e. *Str.1* and *Str.2*); D) projection of community metrics on the same factorial plan. In B) and D) color codes indicate the variables with the most significant contribution to PC1 and PC2. In B) species codes include the first initial of the genus followed by the first three letters of the species names (see the list of species in Fig. 1). In D) S= observed species richness; BM= body mass; CWM= community weighed mean; CWV= community weighed variance; Endo, Epi & Ane= CWM of endogeic, epigeic and anecic scores; Com.1 & Com.2= plot scores on the first two axes of the PCA of earthworm community composition.

The first two axes of the principal component analysis of community composition (i.e. *Com.1* and *Com.2*) explained 35.8% and 16.2% of the total variance of the species abundance table (Fig. A.4A). *Com.1* mostly highlighted an opposition between plots located in the transition zone (1800m and 2000m altitudinal levels with negative scores on PC1), and the rest of the plots (Fig. A.4A). The projection of the species variables on this axis showed that earthworm communities at the ecotone were characterized by the presence of species with high anecic (i.e. *Lumbricus terrestris*, *Aporrectodea longa*) and endogeic scores (i.e. *Aporrectodea caliginosa*, *Octolasium cyaneum*, *Allolobophora chlorotica*) that were almost absent in other altitudinal levels dominated by epigeic species (Fig. A.4B) [3]. *Com.2* was more difficult to interpret and mostly corresponded to an opposition of plots dominated by either *Lumbricus rubellus* or *Lumbricus friendi*, two species with intermediate scores between epigeics and anecics [3] that might be competing and display partially non-overlapping distributions.

The first two axes of the principal component analysis of community structure explained (i.e. *Str.1* and *Str.2*) 37.8% and 17.6% of the total variance of the community table (Fig. A.4C). *Str.1* clearly highlighted the effect of the transition zone, with 1800m and 2000m altitudinal levels having negative scores on PC1 opposed to the rest of the plots (Fig. A.4C). Earthworm communities at the ecotone were characterized by higher earthworm density, biomass and species richness, and by negative scores on the community composition PCA (Fig. A.4D). *Str.2* broadly corresponded to an effect of altitude. Plots located bellow 1800m displayed positive scores on this and were characterized by the presence of species with higher epigeic and anecic scores when compared to higher elevation plots. Only the position of the 2400m level on this axis was unclear, probably because of the low abundance of earthworms that were found at this altitude.

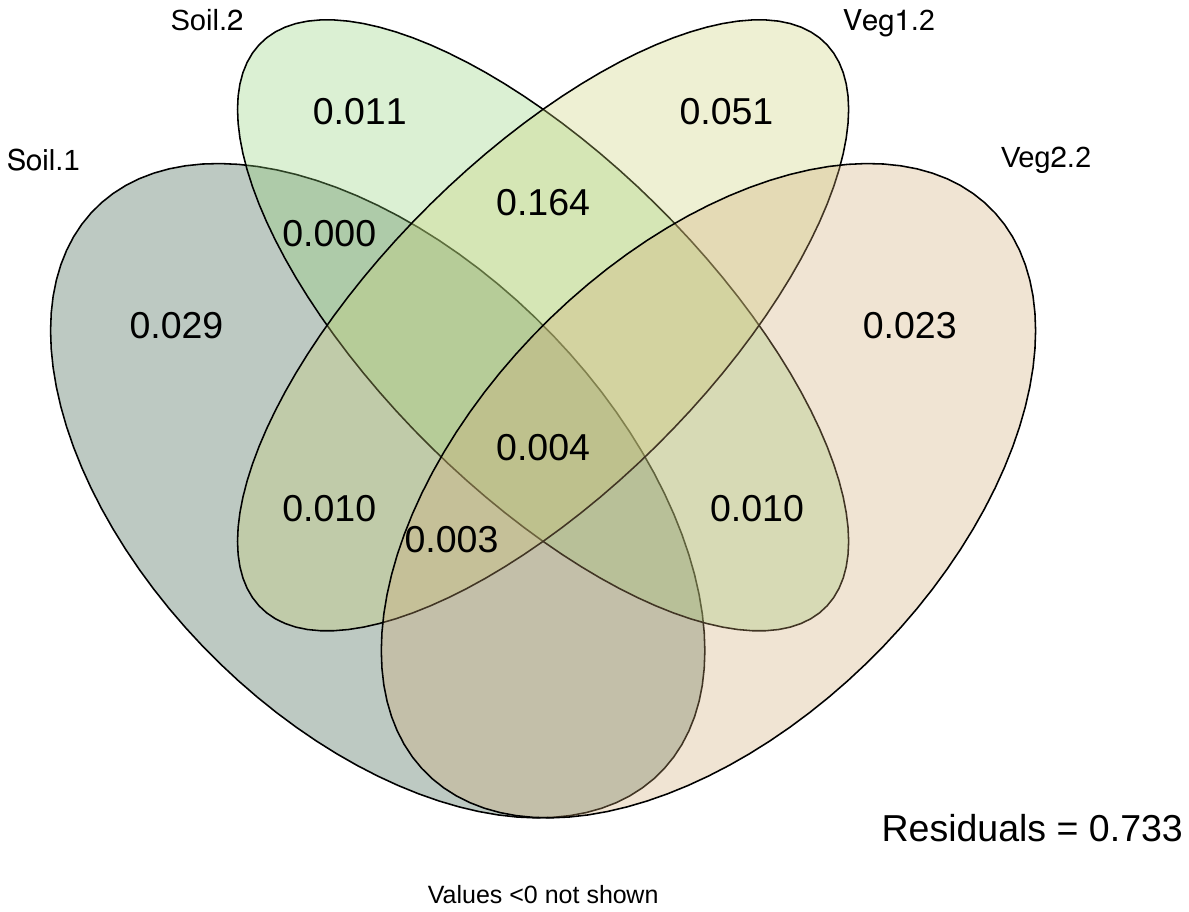

***Figure A.5.*** Results of the variance partitioning analyses highlighting the relative proportion of earthworm community variance explained by the significant synthetic environmental predictors (as identified by the RDA; see Fig. 5), their interactions, and the proportion of the variance not explained by those predictors (i.e. residuals). Synthetic environmental predictors correspond to the plot scores on the first two axes of the PCA of soil properties (*Soil.1* & *Soil.2*), and the second axes of the PCAs of vegetation structure and composition (*Veg1.2* and *Veg2.2*, respectively) as described above.

***References***

[1]	M. Bounous, The influence of plant root systems on soil erodibility and infiltration processes in mountain ecosystems, Masters BEE thesis, University of Montpellier, 2019.

[2]	M.B. Bouché, Lombriciens de France, INRA, Paris, 1972.

[3]	N. Bottinelli, M. Hedde, P. Jouquet, Y. Capowiez, An explicit definition of earthworm ecological categories - Marcel Bouche’s triangle revisited, Geoderma. 372 (2020) 114361. https://doi.org/10.1016/j.geoderma.2020.114361.
